# ERp44 is Required for Endocardial Cushion Development by Regulating VEGFA Secretion in Myocardium

**DOI:** 10.1101/2021.07.02.450976

**Authors:** Youkun Bi, Zhiguang Yang, Meng Jin, Kui Zhai, Jun Wang, Yang Mao, Yang Liu, Mingqin Ding, Huiwen Wang, Fengchao Wang, Guangju Ji

## Abstract

**Rationale:** Endocardial cushions are precursors of the valvoseptal complex that separates the four heart chambers and control blood flow through the heart. Abnormalities in endocardial cushion development lead to atrioventricular septal defects (AVSDs), which affect 1 in 2,100 live births. Several genes have been implicated in the development of endocardial cushions. Specifically, endoplasmic reticulum-resident protein 44 (ERp44) has been found to play a role in the early secretory pathway, but its function in heart development has not been well studied. **Objective:** The goal of this study was to investigate the role of ERp44 in heart development in mice. **Approach and Results:** Using conventional and tissue-specific knockout mouse models, we demonstrated that ERp44 plays a specific role in heart development. ERp44 knockout (KO) mice were smaller in size, and most mice died during early postnatal life. KO hearts exhibited the typical phenotypes of congenital heart diseases, such as abnormal heart shapes as well as severe septal and valvular defects. Similar phenotypes were found in *cTnt-cre+/−; Erp44fl/fl* mice, which indicated that myocardial ERp44 principally controls endocardial cushion formation. Further studies demonstrated that the deletion of ERp44 significantly decreased the proliferation of cushion cells and impaired the endocardial-mesenchymal transition (EndMT), which was followed by endocardial cushion dysplasia. Finally, we found that ERp44 directly bound to VEGFA and controlled its release. **Conclusions:** ERp44 contributes to the development of the endocardial cushion by affecting the EndMT of cushion cells by regulating VEGFA release in myocardial cells.

## 1. Introduction

Many genes have been shown to play essential roles in endocardial cushion development, including molecules involved in transcription, epigenetics, adhesion and migration. These molecules have been summarized by Lin et al^1^. However, given their complexity and importance to regenerative or therapeutic purposes, additional details remain to be elucidated.

ERp44 is a pH-regulated chaperone and belongs to protein disulfide isomerase family^2^. It contains three thioredoxin domains, a, b, and b’, and a flexible carboxy-terminal tail^3^. The CRFS motif in domain a is thought to form a disulfide bridge with ERp44 client proteins^4^. This flexible tail masks the substrate-binding site; thus, the protein is in the off-state at the neutral pH of the endoplasmic reticulum (ER) and in the on-state at the lower pH of the cis-Golgi apparatus, allowing it to capture KDEL receptors^5, 6^. In this manner, ERp44 orchestrates the balance between client protein retention in the ER and their secretion via covalent interactions^7, 8^. Several important secretory factors are reportedly regulated by ERp44, including IgM^9, 10^, adiponectin^11^, SUMF1^12^, and serotonin^13, 14^. Moreover, ERp44 has been implicated in calcium homeostasis. Specifically, ERp44 binds directly to the L3V of inositol 1,4,5-trisphosphate receptor type 1 (IP_3_R1) to inhibit its activity^15^. C160/C212, but not C29, participates in the regulation of IP_3_R1^16^. Huang et al reported that ERp44 may also regulate IP_3_R2^17^. Recently, Wang *et al* reported that ERp44 mutant leads to embryonic lethality using mouse model, but they did not provide a direct reason for this phenomenon^18^. Moreover, Hisatsune *et al* reported that ERp44 regulates blood pressure^19^.

In the present study, we report for the first time that the deletion of ERp44 leads to congenital heart defects; specifically, a loss of ERp44 causes AVC dysplasia arising from the aberrant proliferation of cushion cells and reduced EndMT by directly regulating VEGFA during heart development. Thus, our findings revealed a critical role for ERp44 in endocardial cushion development.

## 2. Methods

### 2.1 Mouse breeding and genotyping

All animal studies were performed in accordance with the relevant guidelines and regulations approved by the Committee on Animal Care of the Institute of Biophysics at the Chinese Academy of Sciences in China. Mice were mated at 5:00 pm and E0.5 was defined as 9:00 am of next day when vaginal plug was detected. Genotyping procedure was performed as previously described^20^.

### 2.2 Generation of ERp44 conventional knockout mice

ERp44 knockout mice were generated by gene targeting the ESCs of the 129 mouse strain and subsequently injecting positive cells into C57BL/6 blastocysts. A Loxp-neomycin-Loxp cassette with a homologous arm was used to replace exon 2 and exon 3 of the *Erp44* gene. The Loxp-neomycin-Loxp cassette was deleted in mutant mice by crossing them with CMV-cre mice. 129×C57BL/6 genetic background mice were backcrossed to C57BL/6 for at least 5 generations. For genotyping, a set of two primers was designed within or without exon 2 and exon 3 for both ERp44-WT-F/R and ERp44-G-F/R. The sequences of the genotyping primers are listed in Table S 1.

### 2.3 Generation of ERp44 conditional knockout mice

The *gRNA* (GTTTTAGAGCTAGAAATAGC) sequence was designed to help Cas9 cut DNA. The sgRNA with T7 promotor was transcribed in vitro using the MEGAshortscript™ Kit (Ambion Life Technologies). Cas9 from Addgene (#41815) including the T7 promotor was transcribed using the mMACHINE T7 ULTRA kit (Life Technologies) and purified using the MEGAclear kit (Life Technologies). The targeting donor was designed to replace exon 2 in the wild-type allele with two flanking loxP sequences and two homologous arms. gRNA, Cas9 and donor DNA were microinjected into C57BL/6 zygotes and transplanted into pseudopregnantcy mice. The founder was genotyped with primers against sequence-F/R, which detected the left loxP sequence, and screen-F/R, which spanned two loxP sequences. The genotype was then verified with Sanger sequencing. The sequences of relative primers were listed in Table S1.

### 2.4 Quantitative real-time PCR

Total RNA was isolated with TRIzol, and cDNA was synthesized with the PrimeScriptTM RT Reagent Kit and gDNA Eraser (TaKaRa). Real-time PCR was performed using the SYBR Green PCR kit(Applied Biological Materials, ABM) and a Rotor-Gene-Q instrument (QIAGEN). The fold-change in target expression between WT and KO mice was calculated using the 2^−ΔΔCt^ method^21^, and GAPDH was used as an internal reference. The primers are shown in Table S 3.

### 2.5 Immunohistochemistry and immunofluorescence

The embryos were excised in cold PBS (10g/L NaCl, 0.25g/L KCl, 1.44g/L Na_2_HPO_4_, 0.25g/L KH_2_PO_4_) and fixed in 4% paraformaldehyde/PBS at 4°C. Gradually dehydrated embryos were carefully embedded in wax under a stereoscope. H&E staining and Alcian blue staining (LEAGENE) were performed according to a standard procedure. Apoptosis was detected with the In Situ Cell Death Detection Kit (Roche) according to the product manual. Images were acquired using a Leica SCN400 Slide Scanner.

For immunofluorescence, frozen sections (8 μm) or adherent cells were fixed (4% paraformaldehyde/PBS) for 10 minutes and antigens on paraffin sections (5 μm) were unmasked with 10 mM sodium citrate buffer. The sections were then washed twice with PBS and permeabilized with 0.3% Triton-100/PBS for 10 minutes before being incubated with blocking solution (ZSGB-BIO) for 1 h at RT. The sections were then incubated with primary antibodies overnight at 4°C. Subsequently, the sections were washed three times and then incubated with secondary antibody for 1 h at RT. Nuclei were counterstained with 1 μg/ml DAPI (Beyotime), and images were acquired using a Leica SP5 confocal microscopy. The following antibodies were used: rabbit anti-Ki67 (1:50, Abcam), mouse anti-SMA (working solution, Abgent), mouse anti-VEGF(1:200, Santa), rabbit anti-ERp44 (1:50, CST), mouse anti-myc(1:50, CWBIO), mouse anti-HA (1:50, CWBIO), goat anti-mouse Alexa Fluor 488 (1:500, Invitrogen) and goat anti-rabbit Alexa Fluor 594 (1:500, Invitrogen). All antibodies were diluted in PBS with 0.3% Triton X-100 and 1% BSA.

### 2.6 AV cushion explant assay

The AV cushion explant assay was modified from previous reports^22^. Briefly, endocardial cushions from the AV cushion were explanted on rat tail collagen gel (BD Biosciences). After an overnight incubation, medium was added (M199, GIBCO; 1% FBS, Invitrogen; 0.1% ITS, GIBCO; 100 U/ml penicillin, 100 mg/ml streptomycin, INALCO), and elongated or spindle-shaped mesenchymal cells were counted after 48 h and analyzed according to Xiong et al^23^. The degree of EndMT of a given explant was assessed based on the number of mesenchymal cells relative to the mean in all control groups, which was defined as 100% EndMT.

### 2.7 DNA constructs

Full-length hVEGFA165 cDNA was amplified from A549 cells and ligated into *pcDNA4* to generate *pcDNA4-hVEGFA165* (VEGFA165) and *pcDNA4-hVEGFA165-myc* (VEGFA165-myc). The ERp44 expression vectors, *pcDNA3.1* (as control), *pcDNA3.1-HA-Erp44* (HA-WT), *pcDNA3.1-HA-C29S* (HA-C29S), and *pcDNA3.1-HA-ΔT*(HA-ΔT) were described previously^16^.

### 2.8 Establishment of cell lines

VEGFA165-myc amplified from *pcDNA4-hVEGFA165-myc* was ligated into PQCXIP vector to generate *pQCXIP-VEGFA165-myc*. Then together with VSV-G and pHIT, they were transfected into Plat E cells for retrovirus packaging. Quality-checked retroviruses infected HEK-293T cells supported by polybrene for follow-up puromycin screening until obtaining modified HEK-293T cell line stably expressing VEGFA165-myc. Details of above procedure referred to previous documents^24, 25^, and it also guided the establishment of H9C2 cell line overexpressing ERp44(OE H9C2).

With regard to generating ERp44 KO H9C2, ERp44-shRNA-F and ERp44-shRNA-R(Table S 2) were annealed, and products were welded into *Lenti-CRISPR-v2* vector digested by *BsmB* I to generate *Lenti-CRISPR-v2-ERp44-shRNA*. This plasmid together with pMD-2G and pAX2 was transfected into HEK-293T cells for lentivirus packaging. Followed by virus infection and puromycin screening, H9C2 cell line with ERp44 KO(KO H9C2) was established.

### 2.9 Transfection, western blotting and co-immunoprecipitation (IP)

HEK-293T cells were cultured in high-glucose Dulbecco’s modified Eagle medium containing 10% fetal bovine serum, 100 U/ml penicillin and 0.1 mg/ml streptomycin (culture medium). The day before transfection, 10^5^ cells were seeded in each well of a 12-well plate. The cells were transfected with Lipofilter (HANBIO). Twelve hours after transfection, the cells were washed twice with PBS, and 2 ml fresh culture medium was added. After 48 h, the culture medium was collected, and 100× protease inhibitor cocktail (AMRESCO) and PMSF (Beyotime) were added. The cells were lysed with RIPA Lysis Buffer containing PMSF. After centrifugation, some of the medium or cell supernatant was mixed with an equal volume of 2× sample buffer with reducing agent (10% β-mercaptoethanol).

For the co-IP experiment, 293T cells grown in 6-well plates were co-transfected with VEGFA165 and ERp44 expression vectors or the VEGFA-myc overexpression vector. After 48-72 h, the cells were washed with PBS and then exposed to 1.2 mM DSP (Pierce) in PBS for 30 min at RT. After quenching with 25 mM Tris-HCl in PBS, the cells were collected in NP40 buffer (PBS containing 100 mM NaCl, 10 mM NEM, 0.5% NP40, 10 mM Tris-HCl 7.4, protease inhibitor cocktail, and PMSF) followed by ultrasonic lysis and centrifugation. The lysates were incubated with anti-HA or anti-myc mouse monoclonal antibody agarose resin (CWBIO) overnight at 4°C. The beads were washed three times with NP40 buffer, and the protein was eluted with 0.2 M glycine (pH 2.5), neutralized with 1.5 M Tris-HCl (pH 9.0). Subsequently, 6x sample buffer with reducing agent was added.

For the western blot analysis, embryonic or neonatal heart tissue was lysed with RIPA Lysis Buffer containing PMSF. The samples were separated by SDS-PAGE and analyzed with anti-ERp44 (CST), anti-VEGFA (Santa Cruz) and anti-β-actin (Proteintech) antibody.

### 3.0 Statistical analysis

Differences were considered significant at p<0.05(*), p<0.01(**) and p<0.001(***) using ANOVA and Student’s t test. All results are expressed as the mean±SEM.

## 3. Results

### 3.1 Generation and characterization of ERp44 knockout mice

To reveal the role of ERp44 in vivo, we generated knockout mice by disrupting the ERp44 gene in embryonic stem cells (ESCs) via homologous recombination (Fig. 1 A). A genotyping study indicated that exon 2 and exon 3 were successfully deleted (Fig. 1 B), and ERp44 expression was undetectable in knockout mice, whereas it was clearly observed in WT mice via western blotting (Fig. 1 C).

**Fig. 1.**
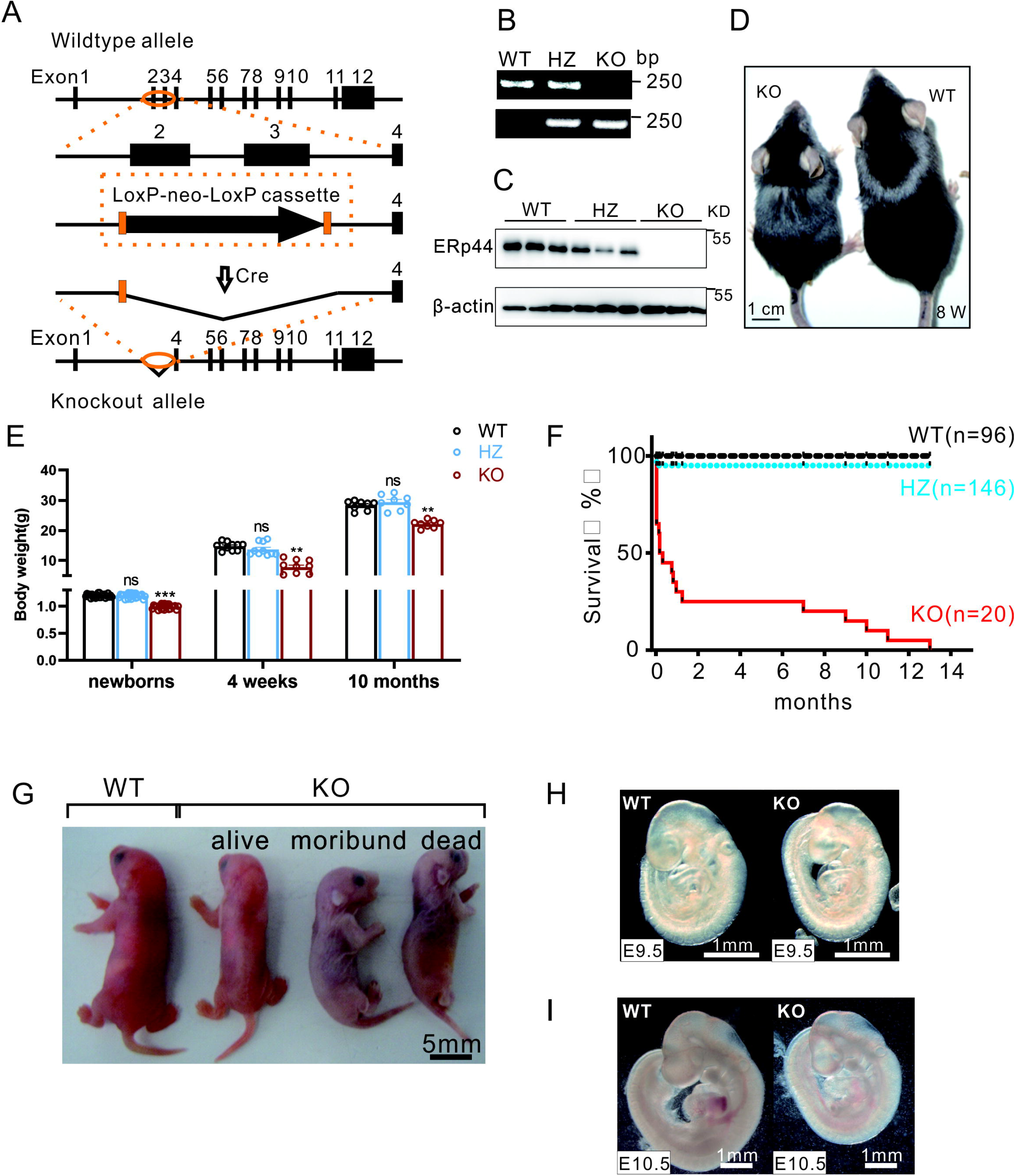
Generation and characterization of ERp44 knockout mice. (A) Strategy used to generate conventional ERp44 knockout mice. (B) Genotyping of WT, heterozygote (HZ) and KO mice. (C) Western blot detection of ERp44 expression in newborn WT, HZ and KO hearts. (D) Photograph of adult WT and KO mice (8 w). (E) Body weights of WT, HZ and KO mice of different ages. (F) Survival curve of WT, HZ and KO mice from the newborn stage to adulthood. (G-I) Photographs of WT and KO mice at postnatal day (G), E9.5 (H), and E10.5 (I).

### 3.2 ERp44 deficiency leads to perinatal embryonic lethality

To assess the effect of ERp44 deletion in mice, we carefully examined the phenotypes of mice. Specifically, ERp44 KO mice were smaller (Fig. 1 D and G) and weighed significantly less (Fig. 1 E) than their littermate controls, and this difference persisted into adulthood. Because the mice were crossed with heterozygotes, the birth rate of KO mice was 21.2% (31/146), slightly lower than the expected Mendelian rate (25%). Most KO mice died within 24 h after birth and exhibited marked cyanosis, and only a few (15% of all KO mice identified after birth) survived to adulthood (Fig. 1 F). Newborn KO mice initially exhibited normal breathing but turned pale (Fig. 1 G) and showed symptoms of tachypnea within a few hours (data not show), suggesting abnormal cardio-pulmonary function. By E9.5, the sizes of KO and WT littermate embryos were similar, but the development of KO embryos was retarded starting at E10.5 (Fig. 1 H and I).

### 3.3 Embryos deficient in ERp44 exhibit cardiac defects

The observed retarded development of the body and high mortality of ERp44 KO mice after birth suggest that cardiac function is developmentally impaired in these mice. To test this hypothesis, we examined the heart under a stereoscope and found that the hearts of newborn KO mice were biventricularly enlarged and exhibited dilated atria (Fig. 2 A and F). A histological analysis of serial sections of the organs of newborn mice stained with hematoxylin and eosin (H&E) demonstrated a high incidence of valvoseptal defects in the hearts of newborn KO mice (Fig. 2 C, F and M), including atrioventricular septal defects (AVSDs), primary aortic septal defects (ASDs) and perimembranous ventricular septal defects (PVSDs). Notably, the muscular septum was unaffected in all KO mice. Pulmonary congestion with focal alveolar edema was also observed in the lungs of KO mice (Fig. 2 G and H), indicating that the pulmonary blood circulation was increased due to a significant left-to-right shunt of the heart. The heart defects of ERp44 KO mice were further confirmed by injecting methylene blue into left ventricle of WT and KO newborn hearts in vivo under a stereoscope. The dye first appeared in the aortic arch in the WT heart, whereas it simultaneously appeared in right ventricle, right atrium and aortic arch in KO mouse hearts (Fig. S 1). We further examined serial sections of the hearts at E14.5-E18.5, when the ventricular septum and primary atrial septum should have been completely closed^26^. Similar with newborn mice, atrioventricular septation dysplasia and dysplastic valve leaflets were observed in E14.5 ERp44 KO hearts (Fig. 2 I, J and M). We also observed a slightly dysplastic valve in a survived adult ERp44 KO mouse (Fig. 2 K and L). These results indicate that ERp44 deletion in mice leads to severe heart defects.

**Fig. 2.**
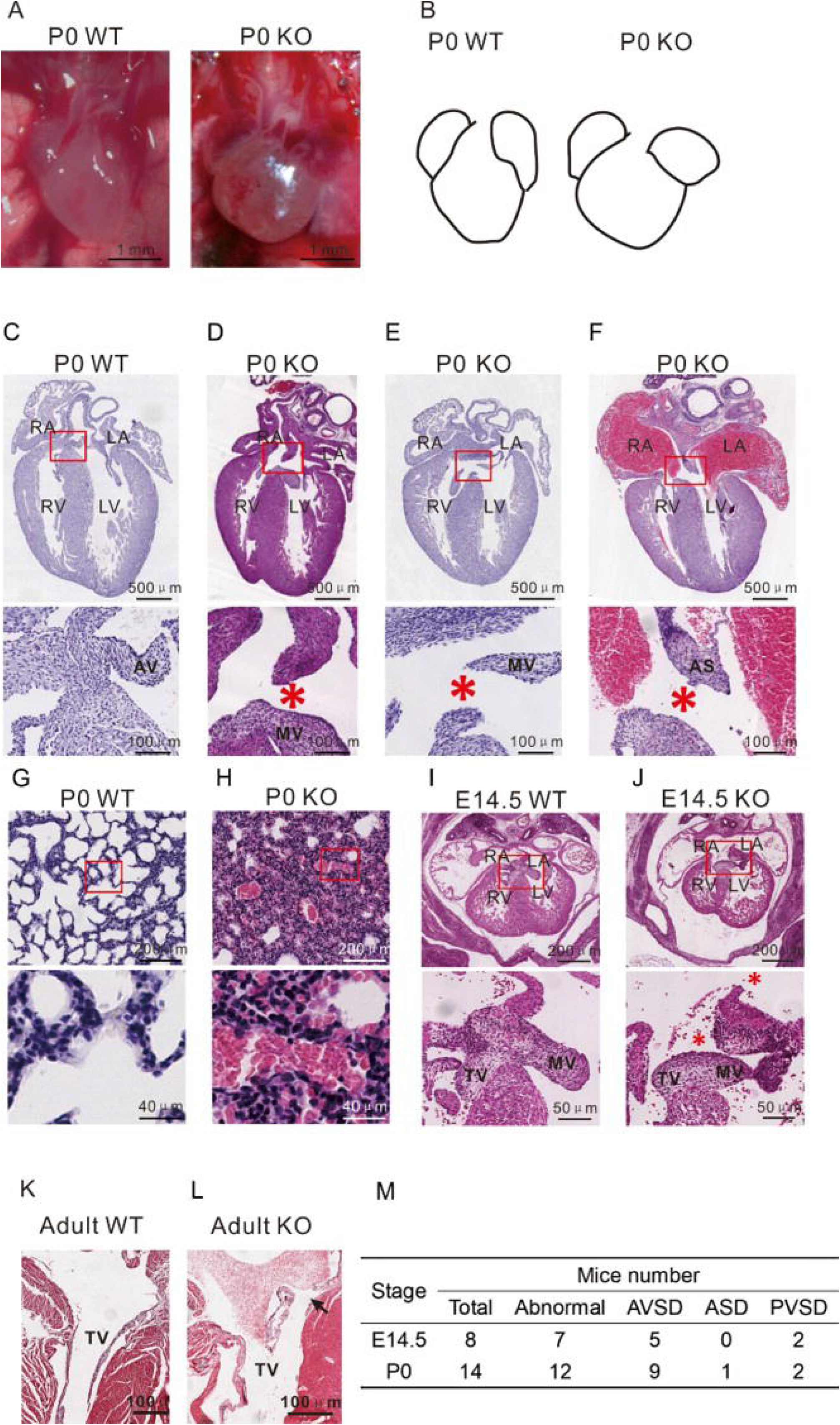
ERp44 deletion causes septal defects in the heart. (A) Photographs of the hearts of WT and KO newborn mice. (B) Schematic map of (A). (C-F) H&E staining of WT (C) and KO (D-F) hearts within 24 h of birth (P0). Asterisks indicate a defective septum. (D) Primary aortic septal defects (PASDs). (E) Perimembranous ventricular septal defects (PVSDs). (F) Atrioventricular septal defects (AVSDs). (G-H) Pulmonary congestion and edema in neonatal WT (G) and KO (H) mice. (I-J) Transverse sections of WT (I) and KO (J) hearts at E14.5. (K-L) Transverse sections of adult WT (K) and KO (L) hearts. (M) Frequency of cardiovascular abnormalities found in KO mice at E14.5 and P0. RA, right atrium; LA, left atrium; RV, right ventricle; LV, left ventricle; AV, aortic valve; MV, mitral valve; AS, atrial septum.

### 3.4 ERp44 deficiency impairs endocardial cushion cell proliferation

Atrioventricular cushions (AVCs) give rise to atrioventricular septation and valves via a series of complex cellular processes^1^. We hypothesized that the heart defects observed in ERp44 KO mice were due to the abnormal development of AVCs. To test this hypothesis, we histologically analyzed both WT and KO embryo hearts at E10.5 and E11.5. The H&E staining of hearts showed fewer cushion cells in the AVCs of KO mice compared with WT mice (Fig. 3 A and B). To explore the reason for this decrease, we examined the proliferation and apoptosis of cushion cells with KI67 and TUNEL assays, respectively. Compared with WT mice, the number of KI67-positive cells was significantly reduced in the AVCs of KO mice (Fig. 3 C and D), whereas the signs of apoptosis were similar in WT and KO AVCs at E10.5 and E11.5 (Fig. 3 E and F). Because impairments of the extracellular matrix (ECM) in cardiac jelly result in the abnormal invasion of endocardial-derived mesenchymal cells^27^, Alcian blue staining was performed at E9.5, which did not show significant differences in acid glycosaminoglycans in the cardiac jelly between WT and KO AVCs (Fig. S 2). These results indicate that the hypoplastic AVCs in ERp44 KO mice are primarily due to a proliferative defect of cushion cells.

**Fig. 3.**
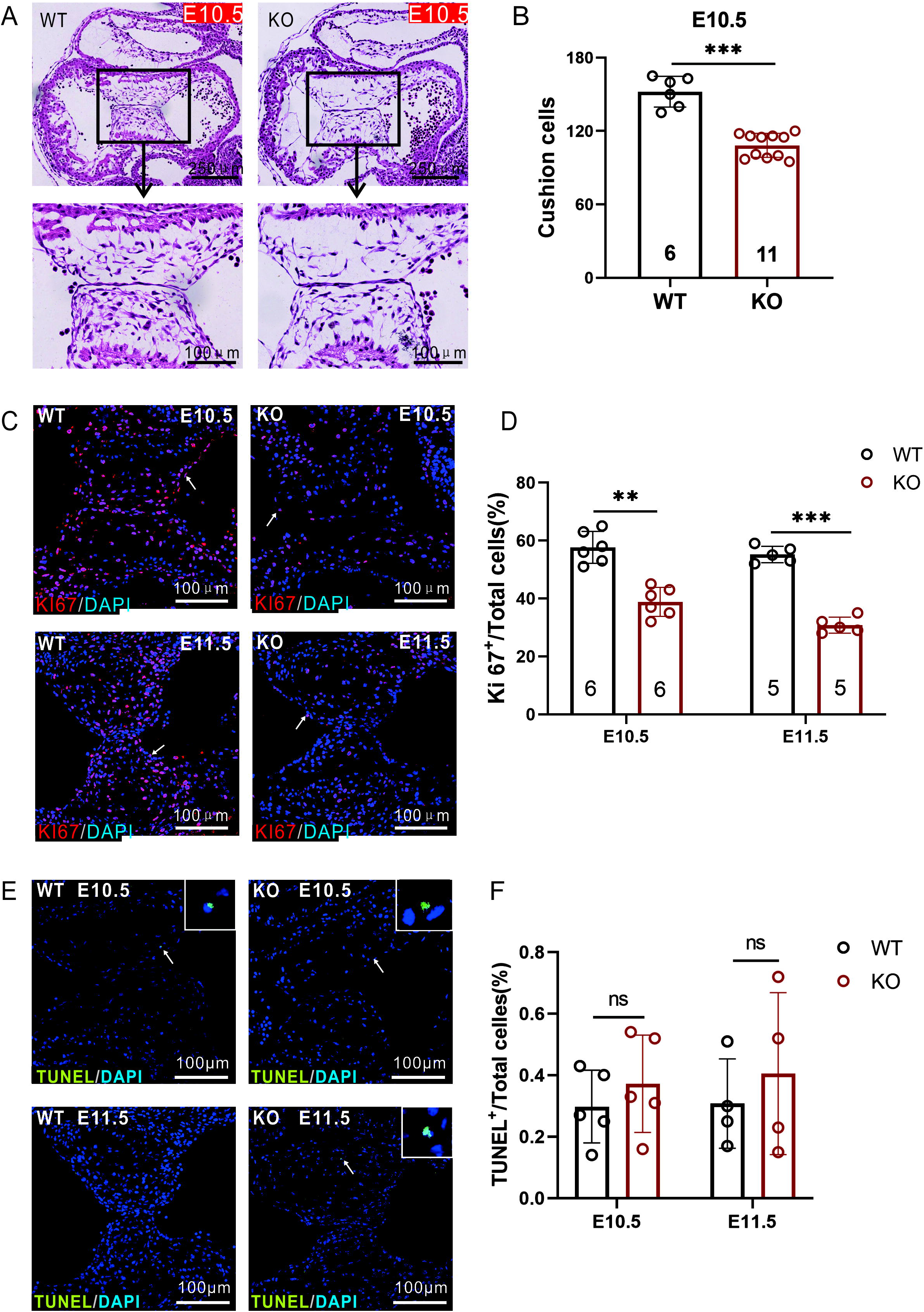
Dysplasia of the endocardial cushion in ERp44 knockout embryos. (A) Histological examination of the atrioventricular canals in WT and KO embryos at E10.5. (B) Quantitation of total endocardial cushion cells. (C and E) Immunostaining of the proliferation marker KI67 (C) and apoptosis marker TUNEL (E) in WT and KO embryos at E10.5 to E11.5. (D and F) Quantitation of positive signals (arrows pointed) in KI67 (D) and TUNEL (F) staining. Arabic numerals in columns represent the experiment number for each genotype.

### 3.5 ERp44 is required for endocardial mesenchymal transition (EndMT)

EndMT is a major cellular process during the development of AVCs^28^. To test whether EndMT is affected in ERp44 KO AVCs, we employed an in vitro assay modified from Feng *et al*^22^. Briefly, AVCs from WT and KO E9.5 were explanted(Fig. 4 A) and cultured on glass plates coated with rat tail collagen gel. After 48 h, large numbers of mesenchymal cells had migrated from the WT explants to the surrounding collagen and exhibited an elongated or spindle-like shape. In contrast, only a few mesenchymal cells were observed around KO explants, and some exhibited an epithelioid-like shape, but the cells from KO AVCs had a similar migratory ability (Fig. 4 B). To confirm our observation, mesenchymal cells were marked with α-SMA antibody and counted, which showed that WT AVCs contained twice as many mesenchymal cells as KO AVCs, as shown in Fig. 4 C and E. These results suggested that ERp44 deletion impairs the transition of mesenchymal cells, but invalidly affect their morphology and migration.

**Fig. 4.**
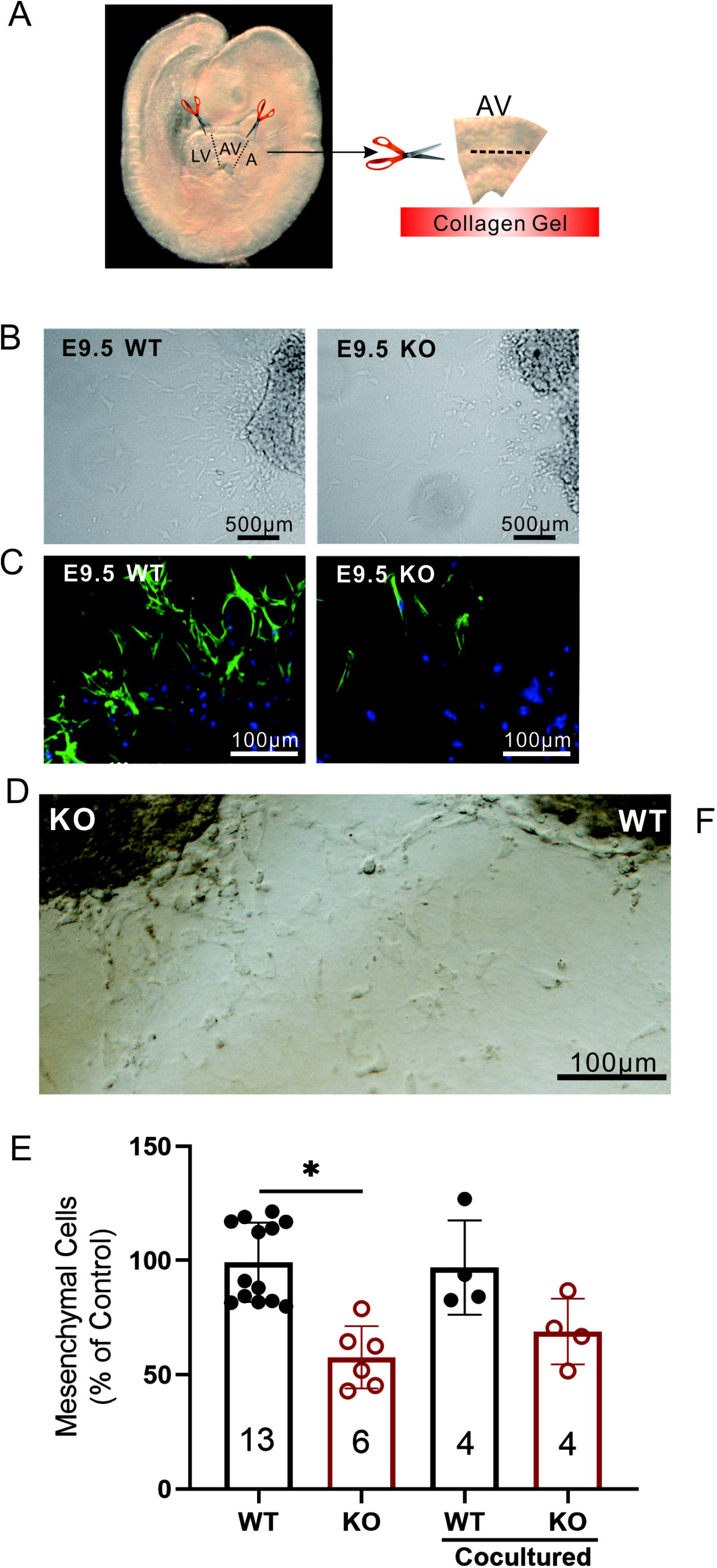
ERp44 is required for EndMT. (A) Schematic diagram to describe the culture of endocardial cushion in vitro. (B) Outgrowths of mesenchymal cells 48 h after endocardial cushion explantation from WT or KO E9.5 embryos as observed by light field microscopy. (C) Immunostaining with antibody for the mesenchymal cell marker anti-αSMA and for nuclei with DAPI from E9.5 explants. (D) EndMT of adjacently cultured WT and KO cushion explants in E9.5 for 48h. (E) Statistics of endocardial cushion explantation trial in (B) and (D).

Furthermore, we investigated whether secreted factors were responsible for the EndMT dysfunction via co-culture trial of WT and KO endocardial cushion in vitro. The atrioventricular septa of WT and KO mice at E9.5 were isolated respectively, and they were adjacently co-cultured in a petri dish coated with rat tail collagen for 48h. Micrographs showed that plenty of spindle-shaped mesenchymal cells migrated from the atrioventricular septum(Fig. 4 D). Co-culture improved the mesenchymal cell migration of KO mice compared with single cultivation. And the mesenchymal migration of WT mice remained unchanged between co-culture and single cultivation(Fig. 4 E). These indicated that co-culture brought out compensation to the mesenchymal cell migration of heart atrioventricular septum in KO mice. Further, we deduced that the lack of some secreted factors, not excessive secretion of inhibitory factors, was responsible for above phenotype.

### 3.6 Myocardial loss of ERp44 is the primary cause of heart defects

EndMT is a complex process in AVCs and is controlled by signals from both the endothelium and myocardium^28^. As shown in Fig. S 3, ERp44 is ubiquitously expressed in the heart. To further assess the autonomous contribution of cells observed in ERp44 knockout mice, ERp44 conditional knockout mice were generated using the CRISPR/Cas9 technique. Briefly, two loxP sequences were inserted between *Erp44* exon 2 in C57BL/6 mice (Fig. S 4A), and the *Erp44* floxed allele (fl) was inactivated in myocardial cells by crossing the mice with *cTnt-cre* mice. *cTNT-cre* delivers Cre within myocardium after E7.5^29^. The hearts and other organs of *cTnt-cre^+/−^;Erp44^fl/+^* mice were collected, and the level of ERp44 mRNA was measured by RT-PCR to confirm deletion. As shown in Fig. S 4 B, ERp44 showed the shortened mRNA mutant band lacking exon 2 (73 bp) in the heart but not in other tissues.

Because *cTnt-cre^+/−^;Erp44^fl/+^* mice were viable and normal, we crossed them with *Erp44^fl/fl^* mice (Fig. S 4 C). Like the conventional KO mice, *cTnt-cre^+/−^;Erp44^fl/fl^* mice were smaller (Fig. 5 A and B) and exhibited fetal lethality, that is, most mice died at birth (Fig. 5 C). Histological analysis of *cTnt-cre^+/−^;Erp44^fl/fl^* embryo hearts at E14.5 and hearts within 24 h of birth(P0) showed AVSDs, ASD and PVSDs (Fig. 5 D to F), which phenocopied the conventional ERp44 KO mice. These results indicate that the myocardial loss of ERp44 is the primary cause of defects.

**Fig. 5.**
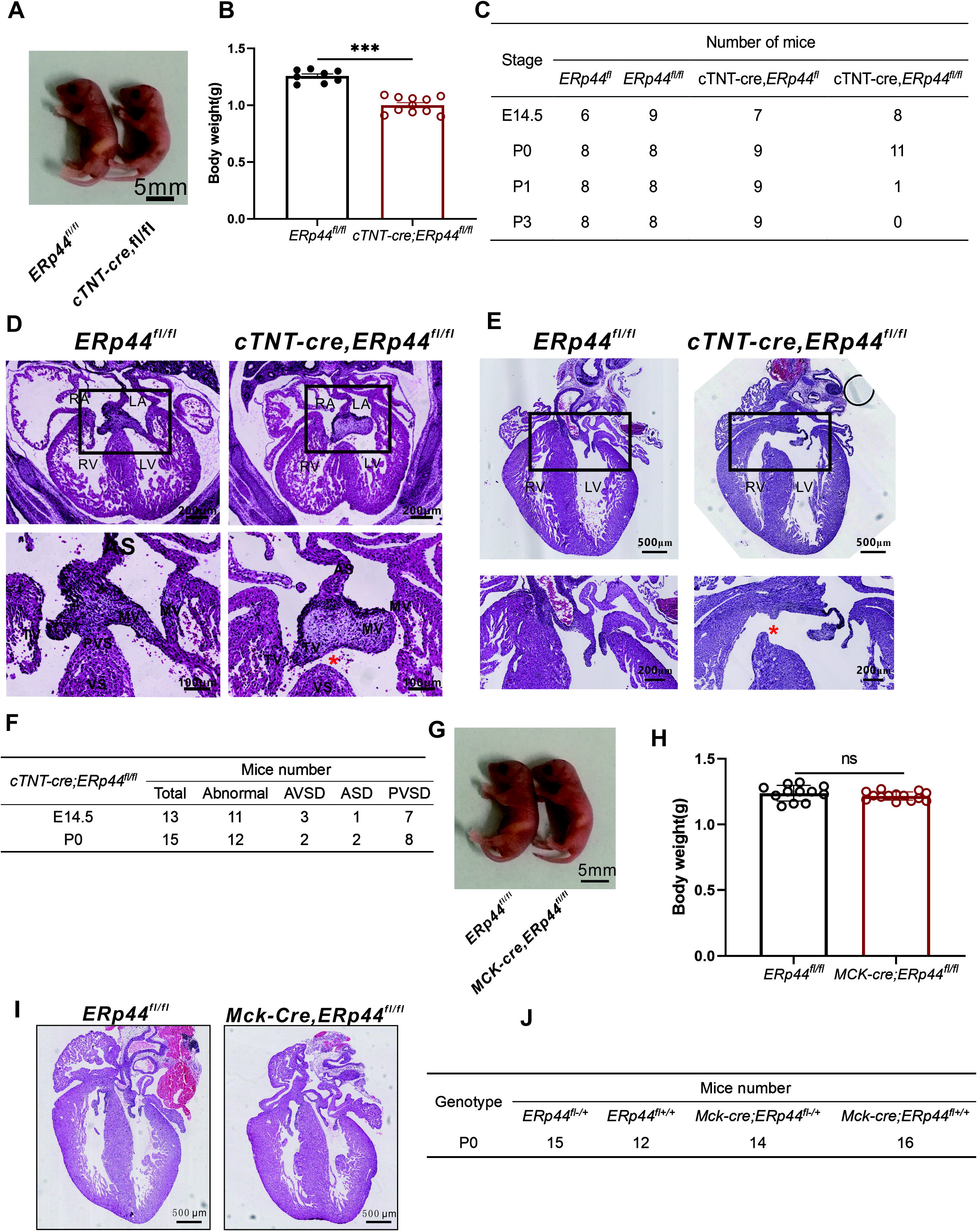
Myocardial-specific deletion of ERp44 leads to AV cushion defects. (A) Photograph of *Erp44^fl/fl^* and *cTnt-cre^+/−^;Erp44^fl/fl^* newborn mice. (B) Quantitation of *Erp44^fl/fl^* and *cTnt-cre^+/−^;Erp44^fl/fl^* body weight. (C) Genotyping results of litters at different stages. (D-E) H&E staining of *Erp44^fl/fl^* and *cTnt-cre^+/−^;Erp44^fl/fl^* at E14.5 (D) and P0 (E). Asterisks indicate defective septum. (F) Frequency of cardiovascular abnormalities found in KO mice at E14.5 and P0. (G) Photograph of *Erp44^fl/fl^* and *Mck-cre^+/−^;Erp44^fl/fl^* newborn mice. (H) Quantitation of *Erp44^fl/fl^* and *Mck-cre^+/−^;Erp44^fl/fl^* body weight. (I) H&E staining of *Erp44^fl/fl^* and *Mck-cre^+/−^;Erp44^fl/fl^* at P0. (J) Genotyping results of litters at P0.

To validate that myocardial loss of ERp44 is important in AVCs development, we crossbred *ERp44^fl^* mice to another muscle-specific Cre tarnsgenic line *Mck-cre. Mck-cre* delivers Cre within myocardium from E13^30^, when AVCs development finishs valve remodeling processes^31^. *We used Mck-cre; ERp44^fl+^* to cross with *ERp44^fl/fl^*. The *Mck-cre; ERp44^fl/fl^* embryos grew normal (Fig. 5 G-H) and did not show obvious congenital heart defects (Fig. 5 I) and the birth rate was normal (Fig. 5 J).

### 3.7 ERp44 KO suppresses VEGFA secretion

Multiple signal pathways affect the development of mouse endocardial cushion, like TGF-β/BMP, Notch, ErbB, Wnt/catenin, and VEGF^32, 33, 34, 35^. We first compared the RNA expression of EndMT signals of the myocardium between E9.5 WT and KO heart AV cushion tissues via RT-qPCR. However, the mRNA level of BMP2/4, TGFβ2/3, Notch1, ErbB3, Wnt3a and VEGF did not significantly differ between these groups(Fig. S 5). Then we detected the protein expression of major EndMT signals, results presented that VEGF expression was significant reduced in KO mice compared with WT(Fig. 6A and B).

**Fig. 6.**
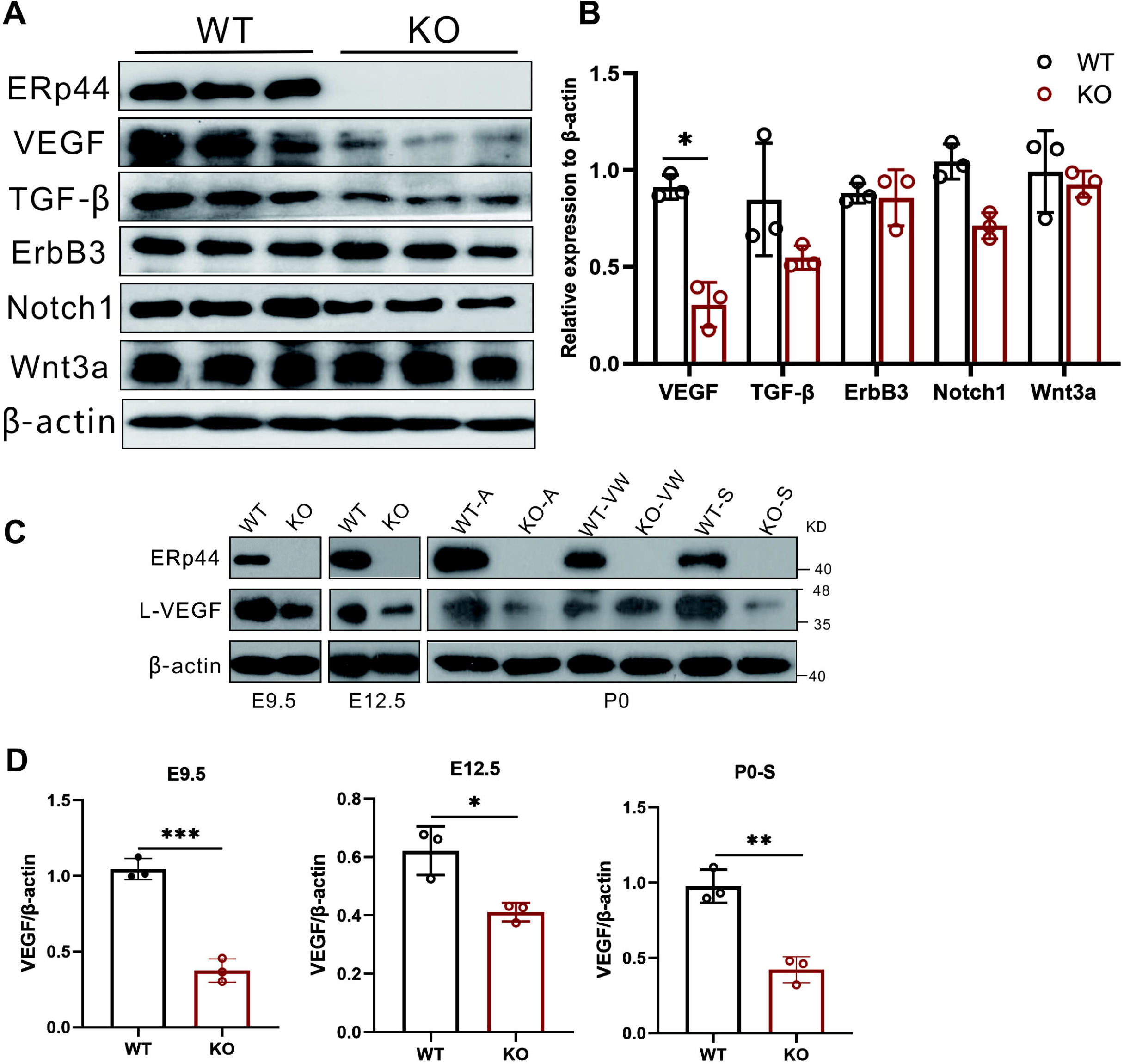
ERp44 KO suppresses VEGFA expression. (A) Western blot detection of multiple crucial proteins that affect the development of mice endocardial cushion targeting WT and KO AVCs tissue in E10.5. β-actin was used as an internal reference. (B) Quantification of protein expression in (A). (C) VEGF detection of WT or KO heart in E9.5, E12.5, and P0, respectively. A, atrium; VW, ventricular wall; S, septum. (D) Quantification of protein expression in (C).

Next, we measured the expression of VEGF at the different development stage of heart, and results indicated that VEGF was all markedly decreased in E9.5, E12.5, and the P0 septal tissues of KO mouse hearts compared with WT mouse hearts (Fig. 6 C and D). Immunohistochemical analysis of heart in E12.5 also supported the downregulation of VEGF in KO mice(Fig. S 6).

To further confirm the relationship between ERp44 and VEGF, we investigated their cytoplasmic localization in H9C2 cells. Micrographs(Fig. 7 A) and Co-IP(Fig. 7 B) analysis revealed ERp44 and VEGF have space colocalization and certain interaction in H9C2. Then, we investigated whether ERp44 contributes VEGF retention within the cell. Four expression plasmids, HA-WT, HA-C29S, HA-ΔT, and Control(pcDNA3.1), were respectively transfected into 293T cell stably expressing VEGF165-myc for 48h to detect the VEGF-myc expression in lysate and supernatant. ERp44 overexpression promoted the VEGF165 expression in cytoplasm, but not in supernatant compared with control. ERp44 mutant(C29S) had no effect on the intracellular and extracellular VEGF165 expression. While truncated ERp44(ΔT) caused the decrease of VEGF165 within cells and supernatant(Fig. 7 C). Unlike ERp44, the C29S mutant did not co-precipitate VEGF, suggesting that efficient binding depends on the Cys29 residue of ERp44(Fig. 7 D).

**Fig. 7.**
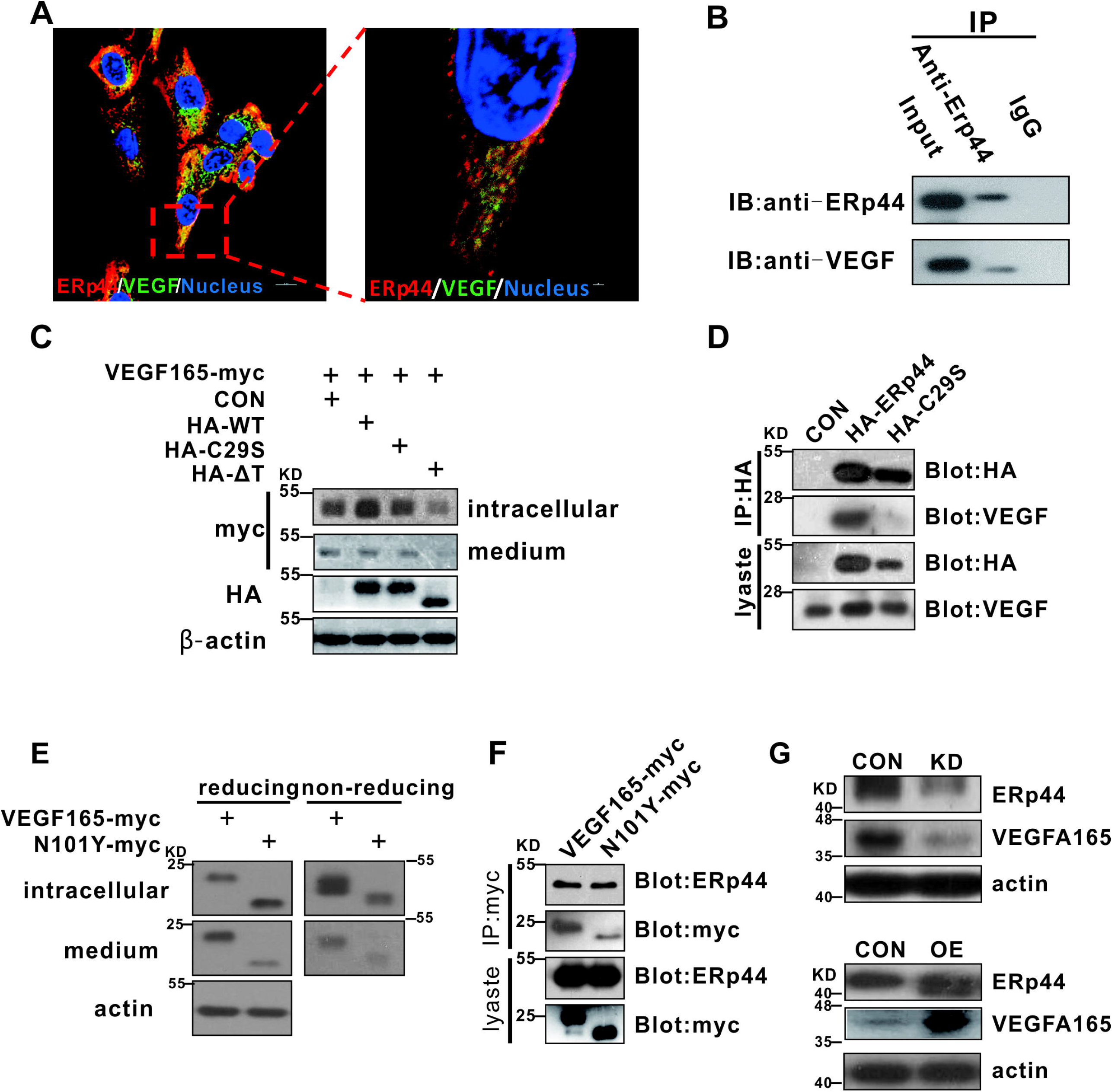
ERp44 affects the development of the endocardial cushion by regulating VEGFA secretion. (A) Micrographs described the colocalization of ERp44 and VEGF in H9C2. The cells were stained with mouse anti-ERp44 (red) and rabbit anti-VEGF (green) antibody. The nuclei were stained with DAPI (blue). (B) Detection of interaction between ERp44 and VEGF in H9C2 by co-immunoprecipitation assay. (C) Detection of intracellular and extracellular VEGF-myc expression in HEK-293T cells stably expressing VEGF-myc after transfecting with ERp44 WT and mutants. β-actin was used as an internal reference. (D) ERp44 interacts with VEGFA, and C29 in ERp44 contributes to the interaction in 293T cells. (E) Glycosylation site N101 of VEGF165 affects its maturation and secretion. (F) Glycosylation site N101 of VEGF165 does not contribute to the interaction of ERp44 and VEGF165. (G) VEGF expression is positively correlated with ERp44 level. OE and KO represent ERp44 overexpression H9C2 and ERp44 knockout H9C2, respectively.

Then we investigated whether glycosylation site N101 of VEGF165 would affect its interaction with ERp44. After transfecting mutant N101 in 293T, Western blot showed VEGF165 monomer decrease and dimer increased within the cell, and both monomer and dimer decreased in the medium(Fig. 7 E). This suggested N101 is crucial for the maturation and secretion of VEGF. Yet, Co-expression of ERp44 with VEGF165 or mutant N101Y revealed that glycosylation site N101 does not affect the interaction of ERp44 and VEGF165(Fig. 7 F). We also detected VEGF level in ERp44 OE H9C2 and ERp44 KO H9C2, respectively. Results showed that VEGF increased in OE H9C2, but notably decreased in KO H9C2(Fig. 7 G). These data suggest that ERp44 is involved in the trafficking of VEGF protein.

## 4. Discussion

Here, we report that ERp44 contributes to the development of endocardial cushion by affecting the proliferation of cushion cells and EndMT. Specifically, myocardial ERp44 principally controlled endocardial cushion formation, and the deletion of ERp44 resulted in heart defects by directly regulating the secretion of VEGFA from the myocardium.

Mice with congenital heart defects were smaller and exhibited growth retardation and embryonic lethality, although some mice may survive after birth, as previously reported^36, 37, 38^. Our findings showed that most KO mice died within the first day of birth, although a few mice survived until adulthood and exhibited slight valve dysplasia and slightly abnormal cardiac morphology (Fig. 1 F and G, Fig. 2 K and L). The few surviving adult KO mice did not show significant cardiac hypertrophy on an echocardiogram (data not shown). Wang *et al* reported that their ERp44 mutant mice died mainly at E12.5^18^, and their surviving adult KO mice showed cardiac hypertrophy, which contradicted our observations. This difference may be due to the fact that they inserted a *LacZ* gene into *Erp44* exon 1 as a fusion protein, which may have caused additional embryonic toxicity, whereas we deleted exon 2 and exon 3 from the *Erp44* allele. Using another ERp44 mutant mouse line (deleting exon 2 and exon 3 of *Erp44*), Hisatsune *et al* recently obtained results similar with ours^19^.

We found that ERp44 KO mice were predominantly AVSD, with some PVSD and ASD and no apparent defect in myocardium. These results showed a diaplasia in the AV cushion development. The proportion of abnormal KO heart was approximately 85%, which was in line with the lethallty rate (about 80%). The fact that CHD phenotype penetrance was lower than 100% was also reported by other studies^38, 39^. Possible explanations for this reason are: 1, the defects were not found by serial section; 2, the mild defects may heal at late developing stages. As no obvious defects were found in KO mice related to OFT cushion development, we did not investigate particularly whether the OFT cushion EndMT was impaired. An explanation of this anomalies is that mesenchymal cells derived from cardiac neural crest cells can migrate in this area to a certain extent^40, 41^.

AV cushion morphogenesis proceeds in the timewindow from E9.5 to E11.5 in mice and has been widely reported to be controlled by signals from both the endocardium and myocardium^28, 42^. After E11.5, heart valve remodeling proceeds^31^. Using myocardial-specific Cre mice (*cTNT-cre* and *Mck-cre*), we observed phenotypes of *cTNT-cre;ERp44^fl/fl^* similar to those observed in conventional knockout mice (Fig. 5 A-F), while *MCK-cre;ERp44^fl/fl^* mice showed no significant phenotype in heart and could grow to adulthood(Fig. 5 G-J). The explanation of this phenomenon is that *cTNT-cre* delivers Cre from E7.5 and *MCK-cre* from E13. These indicates that AV cushion defects are primarily due to hypoplastic EndMT by a loss of myocardial ERp44. Nevertheless, we cannot exclude the role of endocardial ERp44 in our mouse model.

The major signals from the myocardium are BMP2/4, TGFβ2/3, Notch and VEGFA^29, 32, 43^. Although we did not find differences in these genes at the RNA level between WT and KO mice(Fig. S 5), the VEGF decreased in E10.5 ERp44 KO AVCs tissue at the protein level compared with littermate controls(Fig. 6 A and B), and similar phenomenon appeared in different development stages of hearts in KO mice(Fig. 6 C and D), suggesting that less VEGF was secreted in ERp44 KO AVCs.

By using cell line, we confirm that ERp44 interacts with VEGFA(Fig. 7 A and B) and positively controls VEGFA secretion(Fig. 7 G). VEGFA signaling plays a spatio-temporal role in controlling EndMT, and disrupting one VEGFA allele or overexpressing VEGFA causes AV cushion dysplasia^44^. In myocardium, the resting calcium level also regulates EndMT. Specifically, an increase in the intracellular calcium concentration can activate calcineurin, which promotes the translocation of NFATc2/3/4 into the nucleus to regulate the expression of a set of genes, including VEGFA^31, 45^. However, we did not observe differences in the resting Ca^2+^ level in isolated cardiomyocytes between KO and WT mice at P0 (data not shown). The importance of ERp44 in the early secretion pathway has been well studied^9, 10, 19, 46, 47^. Several proteins have been found to interact with ERp44, including IgM^10^, adiponectin^48^, ERAP1^19^, and other clients, sush as VEGFA is speculated to bind to ERp44^49^. Here, we found that ERp44 directly binds to VEGFA and C29 in ERp44 is important for the binding with VEGFA(Fig. 7 C and D). Also, glycosylation site of VEGF N101 is critical for its maturation and secretion(Fig. 7 E), but affects less on the interaction of ERp44 and VEGF165(Fig. 7 F).

In conclusion, we discovered a novel and close link between the ER chaperone protein ERp44 and endocardial cushion defects. Specifically, myocardial ERp44 contributes to the development of the endocardial cushion by affecting the proliferation of cushion cells and EndMT processes, and ERp44 directly regulates VEGFA by binding to the C29 site. The information gained from this study will improve our understanding of the mechanisms underlying AV cushion defects and may also provide a potential diagnostic strategy for congenital heart defects in humans.

## Funding

This work was supported by grants from the National Key Research and Development Project (2019YFA0110402 to GJ) and the National Foundation of Sciences and Technology (31971051, 31771562 to GJ).

## Acknowledgments

The cTnt-cre mouse train was kindly gifted by Prof. Weinian Shou (Department of Pediatrics, Biochemistry & Mol. Biology, Medical & Mol. Genetics, Pharmacology & Toxicology, Indiana University School of Medicine). We thank staff from laboratory animal research center at the Institute of Biophysics for technical assistance and efficient animal care.

## Declaration of interests

The authors declare no competing or financial interests.

## Author contributions

Youkun Bi, Zhiguang Yang, Fengchao Wang and Guangju Ji conceived and designed the experiments. Youkun Bi, Kui Zhai, Meng Jin, Kui Zhai, Jun Wang, Yang Mao, Yang Liu, and Huiwen Wang performed the experiments and contributed reagents/materials/analysis tools. Mingqin Ding and Fengchao Wang performed the design of ERp44 KO mice. Youkun Bi, Zhiguang Yang, and Guangju Ji wrote the paper.

## Supplementary materials

**Fig. S 1.** Exploration of hemodynamics in vivo. Methylene blue (1mg/ml) was gently injected into the left ventricle of newborn WT or KO hearts (0.00 s). The dye passed through the aortic arches in both hearts (0.33 s) and filled the right atrium and auricle of KO mice 1.67 s later instead of passing into systemic circulation, similar with WT mice (2.00 s).

**Fig. S 2.** Alcian blue staining of AVCs in E9.5 WT and KO embryos.

**Fig. S 3.** ERp44 expression in embryonic heart. (A and B) Whole-mount immunohistochemistry at E9.5. Sagittal-view of AVC (A) and OFTC (B) were shown. (C and D) ERp44 immunohistochemistry at E11.5 (C) and E15.5 (D). (C) was vertical section and (D) was transverse sections. AVC, atrioventricular cushions; OFTC, outflow tract cushion; RA, right atrium; LA, left atrium; RV, right ventricle; LV, left ventricle; AV, aortic valve; MV, mitral valve; AS, atrial septum.

**Fig. S 4.** Strategy used to generate conventional ERp44 knockout mice. (A) gRNA and Cas9 mRNA were injected into C59BL zygotes. The targeting donor was designed to replace exon 2 of the wild-type allele with two flanking loxP sequences (shown LoxP). Two homologous arms were used for homologous repair. (B) Schematic showing the ERp44 deletion in myocardial cells by *cTnt-cre*. RT-PCR showed an additional 73-bp-shorter band in the heart but not liver of *cTnt-cre;Erp44^fl/+^mice*. (C) Schematic showing the crossing strategy. *cTnt-cre^+/−^;Erp44^fl/^* mice were crossed with *Erp44^fl/fl^* mice.

**Fig. S 5.** Total RNA was harvested from E9.5 WT (n=3) and KO (n=3) AV cushion tissue, and qPCR was performed to measure WT and KO gene expression as indicated. *P<0.05.

**Fig. S 6.** Immunohistochemical analysis of VEGF expression in E12.5 embryonic heart.

## Notes

### Competing Interest Statement

The authors have declared no competing interest.

